# Optimization-based synthesis of stochastic biocircuits with statistical specifications

**DOI:** 10.1101/187823

**Authors:** Yuta Sakurai, Yutaka Hori

## Abstract

Model-guided design has become a standard approach to engineering biomolecular circuits in current synthetic biology. However, the stochastic nature of biomolecular reactions is often overlooked in the design process. As a result, cell-cell heterogeneity causes unexpected deviation of biocircuit behaviors from model predictions and requires additional iterations of design-build-test cycles. To enhance the design process of stochastic biocircuits, this paper presents a computational framework to systematically specify the level of intrinsic noise using well-defined metrics of statistics and design highly heterogeneous biocircuits based on the specifications. Specifically, we use descriptive statistics of population distributions as an intuitive specification language of stochastic biocircuits and develop an optimization based computational tool that explores parameter configurations satisfying design requirements. Sensitivity analysis methods are also developed to ensure the robustness of a biocircuit design. These design tools are formulated using convex optimization programs to enable efficient and rigorous quantification of the statistics without approximation, and thus, they are amenable to the synthesis of stochastic biocircuits that require high reliability. We demonstrate these features by designing a stochastic negative feedback biocircuit that satisfies multiple statistical constraints. In particular, we use a rigorously quantified parameter map of feasible design space to perform in-depth study of noise propagation and regulation in negative feedback pathways.

## Introduction

The last two decades of intense efforts in synthetic biology have greatly expanded our ability to build synthetic biomolecular circuits by adopting many concepts and techniques from engineering disciplines. Model-guided design is one of such examples that have been routinely used to create safe and robust control systems in traditional engineering [1] and have been adopted in the design process of biocircuits [2]. To date, many biocircuit modules were engineered with the help of model-based simulations, including logic gates [3, 4, 5], oscillators [6, 7, 8, 9] and genetic memory [10, 11] to name a few. A current challenge of biocircuit engineering is to integrate these circuit modules and build systems for complex operations in the real-world environments, which requires more stringent reliability of each circuit module. As is the case with any engineering systems, a first key step to the robust design of such complex systems is to set appropriate and well-defined performance norms that can specify all the necessary features of systems' behavior. Mathematical and computational tools then facilitate design space exploration to nd parameter con gurations that achieve pre-specified performance requirements. Compared with this ideal, the current design process of biocircuits is still far immature in that models are used mostly for simulations to gain only qualitative insights rather than for quantitatively guaranteeing the performance of biocircuits by fully bene tting from advanced theory and algorithms. This motivates us to develop computational frameworks that streamline the design process by systematically certifying and optimizing the performance levels of biocircuits.

One of the important features that should be carefully considered in the design process of biocircuits is cellular heterogeneity. In biological cells, the low copy nature of molecules induces randomness of molecular collision events that fire chemical reactions, resulting in the large variation of biocircuit states across cell populations even if the cells are genetically identical and grown in the same condition [12, 13, 14]. In many cases, the signal-to-noise ratio of biocircuits is much lower than that of mechanically and electronically engineered systems. Although noise attenuation has been a rule of thumb in engineering, recent studies demonstrated opposite strategies to take advantage of the highly stochastic nature of biomolecular reactions and design biocircuits that operate collectively at a population level [15, 16, 17, 18]. For example, collections of binary outputs from stochastic biocircuits can form a graded response that enables analog decision making in highly stochastic environments [18, 19]. These examples illustrate that we can design novel mechanisms that are different from those in traditional systems to control biocircuits by actively leveraging the heterogeneous responses of cell populations.

To enhance the design process of stochastic biocircuits, the first key step is to produce specifications that can capture all the necessary design features using simple but well-defined performance metrics. For this purpose, useful criteria would be descriptive statistics such as covariance, correlation, the coefficient of variation (CV) and Fano factor in addition to the population mean values. In current synthetic biology, the majority of studies uses Monte Carlo based stimulations of single cell trajectories to approximately evaluate these statistics of population distributions[20]. To complement the time-consuming nature of the Monte Carlo approach, other computational tools are available to directly quantify population distributions [21, 22] and raw moments [23, 24, 25, 26] without running simulations. However, these methods are not designed to directly compute descriptive statistics of biocircuits, which makes it difficult to further develop systematic design tools that can handle statistical biocircuit specifications.

In this paper, we present a design-oriented computational framework that directly calculates steady state statistics of stochastic biocircuits and their sensitivity to parameter perturbations (Fig. 1). Building upon a moment computation approach [26, 27, 28], we formulate the biocircuit design problems in the form of convex optimization programs [29], which enable efficient evaluation of the statistics and its sensitivity without running time-consuming simulations of single-cell trajectories. Our optimization based synthesis approach is capable of characterizing feasible design space that satis es multiple and possibly incompatible performance specifications with mathematical rigor. Thus, it greatly facilitates rational engineering process of noisy biomolecular reactions. In addition, the sensitivity analysis ensures robustness against parameter uncertainty by compensating for errors due to model misidenti cation and perturbations to the host cell environments.

We use the proposed algorithms to explore the design parameter space of a self-negative feedback biocircuit, where a repressor protein regulates its own expression. Specifically, we run the convex optimization programs and obtain a parameter map of the feasible design space with which the negative feedback biocircuit satis es pre-specified performance requirements. Interestingly, the parameter maps indicate the existence of an optimal translation rate that minimizes the CV of the repressor copy numbers, implying that increasing the copy number of the protein by strong translation does not necessarily attenuate the noise due to some effects of negative feedback pathways. To better understand the mechanisms, we perform in-depth study of the propagation of the noise and identify two sources of noise in a trade-off relation, which produces an optimal con guration that minimizes the CV of the repressor protein. Informed by these analyses, we determine a design strategy of the negative feedback biocircuits. Sensitivity analysis is further performed to assess the robustness of our design against parameter perturbations.

**Figure 1:**
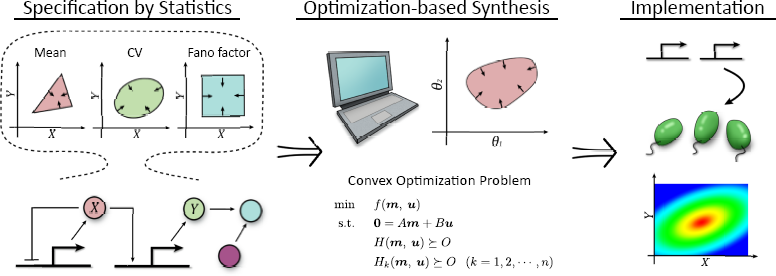
Overview of the optimization based synthesis approach. The optimization program allows for rigorous characterization of parameter space that satis es given statistical specifications.

## Results and Discussion

### Mathematical model of stochastic biocircuits

We start with a general model of stochastic biomolecular reactions and introduce an ordinary differential equation (ODE) model that describes the evolution of stochastic moments of biocircuits. Suppose a biocircuit consists of *n* species of molecules that vary in time and *r* types of chemical reactions. The copy numbers of the *n* molecules, or the state of bio-circuits, uctuate randomly in time and become heterogeneous between cells due to the stochastic chemical reactions. To model the stochastic dynamics, we denote the copy number of the *i*-th molecule by *x_i_* (*i* = 1, 2, …, *n*) and define the probability that there are *x* = [*x*_1_, *x*_2_, …, *x_n_*]^*T*^ molecules in a cell at time *t* by *P_x_*(*t*). As an illustration example, we consider a simple transcription-translation process in Fig. 2A. In this example, mRNA and protein copy numbers are the state variables of the biocircuit (*n* = 2), and there are four reactions (*r* = 4), transcription, translation and degradation of mRNA and protein. The heterogeneity of a cell population is then captured by the joint distribution of mRNA and protein copy numbers 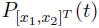, where *x*_1_ and *x*_2_ denote the copy numbers of mRNA and protein, respectively.

**Figure 2:**
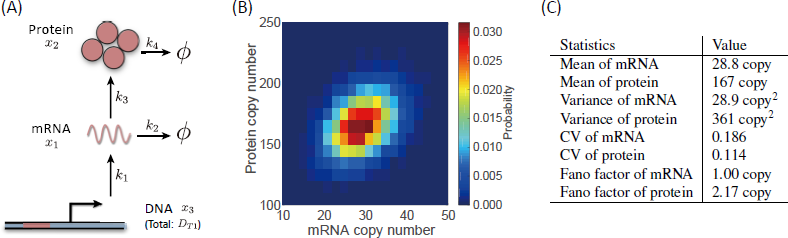
(A) Schematic diagram of a simple transcription-translation biocircuit. (B) Joint population distribution of mRNA and protein copy numbers at *t* = 1440 min. (C) Summary statistics of the population distribution.

In general, the dynamics of the probability distribution *P_x_*(*t*) are modeled by a set of ODEs called Chemical Master Equation (CME) [30].

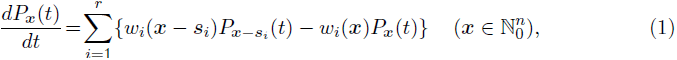

where *w_i_*(·) is a propensity function (reaction rate) associated with the *i*-th chemical reaction (*i* = 1, 2, …, *r*), and *s_i_* is a *n*-dimensional row vector representing the stoichiometry of the *i*-th reaction. We assume that the reactions are elementary, and thus *w_i_*(·) is a polynomial of *x_i_* (*i* = 1, 2, …, *n*) [31]. Specific forms of *w_i_*(·) and *s_i_* for the transcription-translation process in Fig. 2A are summarized in Supporting Information S.2. It should be noted that the entries of the vector *x* in (1) take all combinations of nonnegative numbers, and thus, the CME is composed of in nitely many coupled equations. Although it is hard to analytically solve the equation in terms of *P_x_*(*t*), it is possible to simulate many numbers of single-cell trajectories using a Monte Carlo approach [20] and obtain approximate distributions of *P_x_*(*t*) as illustrated in Fig. 2B. To quantitatively capture the important features of population distributions, widely used statistics are the mean 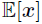 and the covariance 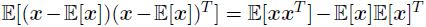, which are the first two central moments of the distribution. Other examples of useful descriptive statistics are the coefficient of variation (CV) 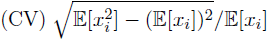 and Fano factor 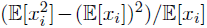, which quantify the dispersion of distributions. In particular, Fano factor become exactly one if the distribution *P_x_*(*t*) is Poisson. Correlation 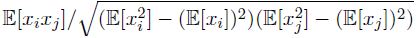, is also a useful measure when we are interested in the relation between two molecules.

The design-oriented computational framework presented in this paper complements the Monte Carlo approach by allowing for rigorous evaluation of descriptive statistics without approximation. For this purpose, we first derive an ODE model that describes the dynamics of raw moments, or a moment equation for short, based on the CME (1). To elucidate the following mathematical development, we first consider a specific model for the transcription-translation process in Fig. 2A. Let *m* denote a vector of raw moments 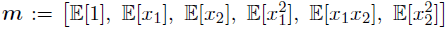, and consider to derive an ODE model for ***m***. The basic idea for the derivation is to multiply 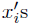 to both sides of the CME (1) and take the sum of 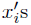 for all nonnegative numbers (Supporting Information S.2 for details). Using this approach, we obtain the moment dynamics as

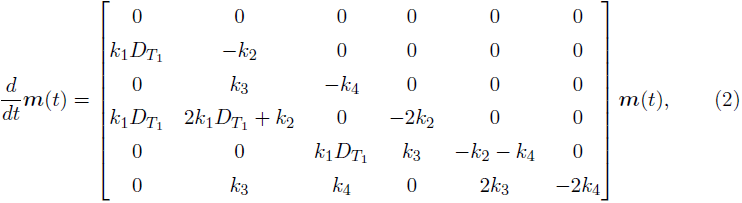

where the rst entry of 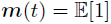 represents the sum of the zero-th order moments of *x*_1_ and *x*_2_, guarantees the sum of the probability *P_x_*(*t*) to be one. Note that no approximation is used in the derivation. In particular, the equation (2) is a linear ODE, and thus, we can rigorously calculate the raw moments ***m*** by solving (2).

When the reactions reach steady state, the left-hand side of (2), which is the time derivative of ***m***(*t*), goes to zero, leading to a set of linear equations. Thus, we solve the linear equations to obtain the first and second order steady state raw moments of the protein copy number, *x*_2_ as

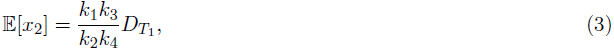

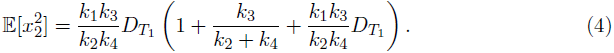

Using these solutions, we can further compute the variance of the protein copy number as

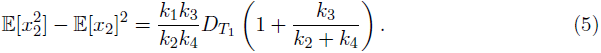

Substituting parameter values, we con rm that the analytic solution indeed agrees with the simulated statistics (Fig. 2C). In the design process of stochastic biocircuits, these analytic solutions are useful for characterizing the parameter space that satis es design requirements and narrowing possible combinations of genetic parts of biocircuits.

### Computing descriptive statistics using semi-algebraic optimization

Unfortunately, analytic solutions are not necessarily available when a biocircuit of interest is slightly more complicated since, in general, a moment is dependent on other (higher order) moments, and the order of a moment equation is in nite. More formally, we define raw moments of a distribution *P_x_*(*t*) by

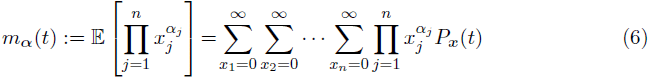

with 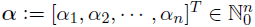 and refer to the sum 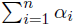 as the order of the moment. A general form of the moment equation is then obtained as

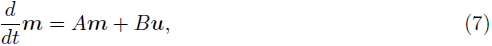

where *A* and *B* are constant matrices, ***m*** is a vector of raw moments up to the *μ*-th order, and ***u*** is a vector of the *μ* + 1-th or higher order moments ***u*** (see Supporting Information S.3 for details). Note that (2) is a special case of (7) with *B* = **0**. Equation (7) implies that the *μ*-th order moments ***m***, which are the moments of our interest, depend on the higher order moments ***u***. Thus, it is not possible to uniquely determine the solution of the steady state moment equation *A****m*** + *B****u*** = **0** since there are more variables than equations. In fact, analytic steady state moments are available only in the special case of *B* = **0**, in which case ***m*** is obtained by solving *A****m*** = **0** as shown in (3) and (4). In general, *B* = **0** holds if and only if all reactions are the zero-th or the first order, that is, the reaction rates *w_i_*(***x***) are affine in ***x*** (see Supplementary Information S.3).

To see an example, we consider a negative feedback biocircuit in Fig. 3A, where the expression of the repressor protein is self-regulated by the negative feedback. This biocircuit, despite a slight extension of Fig. 2A, contains a bimolecular reaction, namely the binding of the repressor to the promoter whose propensity function is given by *w*(***x***) = *k*_5_*x*_2_*x*_3_. As a result, the matrix *B* in (7) is no longer zero, and there are in nitely many solutions for the steady state moment equation unless we know additional information that links ***m*** and higher order moments ***u***.

To constrain the solution, a key observation is that the variables ***m*** and ***u*** must constitute moments of some probability distribution defined on the positive orthant {[***x***_1_, ***x***_2_, …, *x_n_*]| *x_i_* > 0, *i* = 1, 2, …, *n*}. An obvious necessary condition is that all entries of ***m*** and ***u*** must be positive according to the definition (6). In fact, there are tighter conditions that the variables ***m*** and ***u*** must satisfy to be moments of some probability distribution (Proposition A.1 in Supporting Information)[32, 33]. Incorporating these conditions, we can narrow possible combinations of raw moments ***m*** and ***u*** as specified by the following proposition.

**Proposition**. Consider stochastic chemical reactions modeled by the CME (1). The steady state moments of the probability distribution satisfy

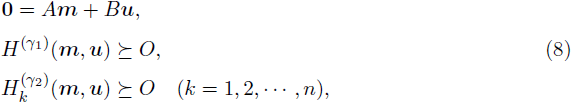

where the matrices 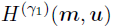 and 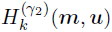 represent moment matrices defined in (A.21) and (A.22) of Supporting Information. The symbol 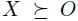 represents that a matrix ***x*** is positive semidefinite.

This proposition implies that the raw moments ***m*** and ***u*** lie in the semi-algebraic set specified by (8). Although it is hard to uniquely determine ***m*** and ***u*** from these conditions, equations (8) imply that we can computationally search for possible combinations of ***m*** and ***u*** based on (8). In particular, we can find the upper and/or lower bounds of statistical values such as the covariance, CV and Fano factor of molecular copy numbers. In what follows, we show that the problem of finding the upper and the lower bounds of these statistics can be recast as a mathematical optimization problem, which we can solve efficiently using existing algorithms of mathematical programming.

We consider the negative feedback biocircuit in Fig. 3A and define ***x***_1_, ***x***_2_ and ***x***_3_ as the copy numbers of mRNA, repressor protein and free DNA, respectively. Our goal here is to compute the mean and the CV of the repressor copy number ***x***_2_ without running stochastic simulations of single-cell trajectories. To this end, we use the semi-algebraic constraint (8) and formulate a maximization problem of the mean and the CV as

**Figure 3:**
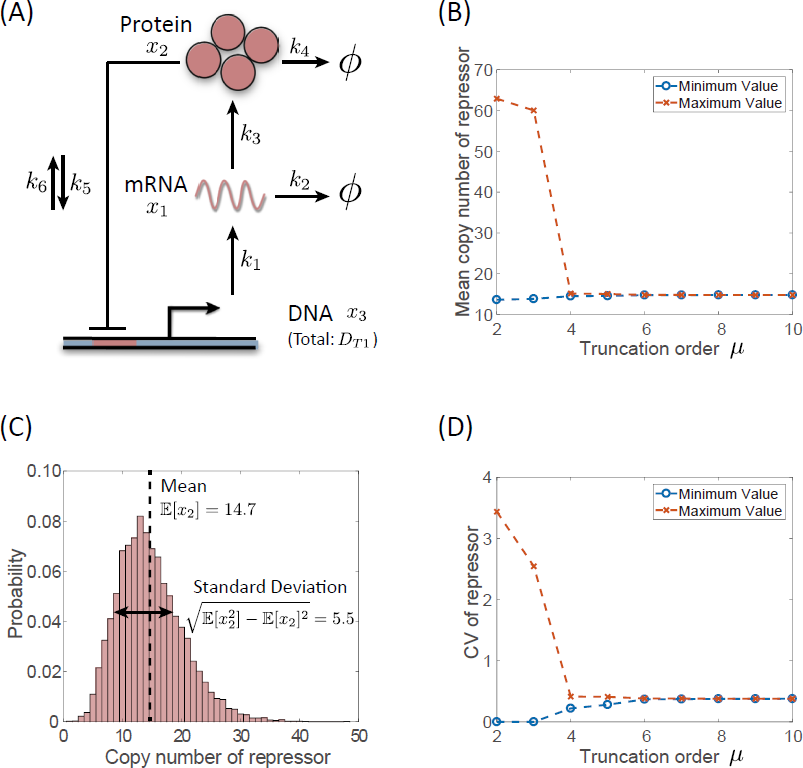
(A) Schematic diagram of a negative feedback biocircuit. (B) Distribution of the repressor copy number at *t* = 1440 min. (C) Computed upper and lower bounds of the mean copy number for different truncation orders *μ*. (D) Computed upper and lower bounds of the variance of the copy number for different truncation orders μ.

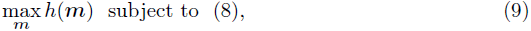

where *h*(***m***) is defined by 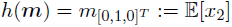 for the mean, and 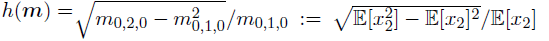 for the CV, respectively. Then, the solution of this problem gives upper bounds of the mean and the CV, respectively. An advantage of using the optimization approach is that we can leverage efficient algorithms for mathematical optimization, whose techniques were extensively studied in engineering science. In particular, the computation of these summary statistics can be recast as semi-definite programming (SDP) [34, 29], which is a subclass of convex optimization program with many practically useful properties such that it allows for finding global minimum (or maximum) with much less computational efforts than other mathematical optimization (Supplementary Information S.4 for details).

Using the SDP approach, we computed the lower and the upper bounds of the mean repressor copy number, 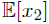 for different values of *μ*, which is a user-specified parameter that determines the largest order of moments in the vector ***m*** in (8) (Fig. 3B). Note that the lower bounds are obtained by solving a similar form of optimization that maximizes −*h*(***m***). As we increase *μ*, the gap between the upper and lower bounds decreases in general, allowing for better estimation of statistical values at the expense of computational time (see Supporting Information). For the biocircuit in Fig. 3A, the estimated mean copy number of the repressor 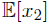 converged to 14.7 (Fig. 3B), which agrees with the mean value of the approximate distribution computed by Monte Carlo simulations [20] (Fig. 3C).

The upper bound of the CV is an important performance norm to quantify the dispersion of the population distribution. Since the optimization of the CV in (9) is not directly solvable by SDP, we developed a procedure to recast the optimization problem (9) into a SDP form by introducing additional variables (see Supplementary Information S.4). Using this approach, we computed the upper bounds of the CV for different values of *μ* as illustrated in Fig. 3D, where the lower bounds were also computed for a reference. We observe that, similar to the mean value, the lower and the upper bounds of the CV approach as we increase the order of the moments *μ*, which implies that the estimation becomes more accurate.

**Table 1:**
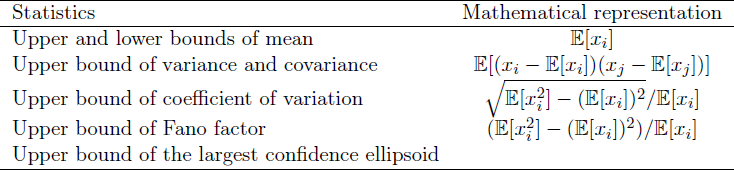
Summary statistics of stochastic biocircuits computable by semidefinite programming

Similar mathematical techniques apply to other descriptive statistics and allow us to rigorously compute covariance, Fano factor and con dence ellipsoids of the molecular copy numbers using semidefinite programming (Table 1). Specific forms of these optimizations and their mathematical proofs are summarized in Supplementary Information S.4. As an illustrative example, we computed the largest con dence ellipsoid of mRNA and protein copy numbers of the transcription-translation circuit in Fig. 2A. The two-dimensional con dence ellipsoid allows for visualizing the correlation between the two molecules (Fig. S.1). In the design process, the con dence ellipsoids would be useful to investigate how tightly a target molecule is regulated by an upstream molecule. Other optimizations will be demonstrated in the following sections along with the design examples of a negative feedback biocircuit.

### Synthesizing biocircuits with statistical design specifications

The process of biocircuit engineering requires many iterations of design-build-test cycles to achieve prescribed performance requirements. Since the specifications are possibly incompatible or con icting, computational design tools are important to efficiently explore and find the feasible design space of biocircuits. Our optimization approach allows for rigorous characterization of biocircuit parameter space satisfying multiple design requirements described by statistical constraints (Table 1). Specifically, we use a set of inequalities to mathematically specify the design requirements of biocircuits. For example, 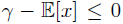 implies that the copy number of a molecule, say ***x***, must be more than *γ*, and 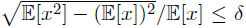 implies that the CV must be less than *δ* at steady state. More formally, we denote biocircuit specifications by *f_i_*(***m***) ≤ 0 (*i* = 1, 2, …, *s*) with raw moments ***m***. Using the convex optimization presented in the previous section, we can rigorously determine whether a given circuit design satis es these performance specifications.

**Figure 4:**
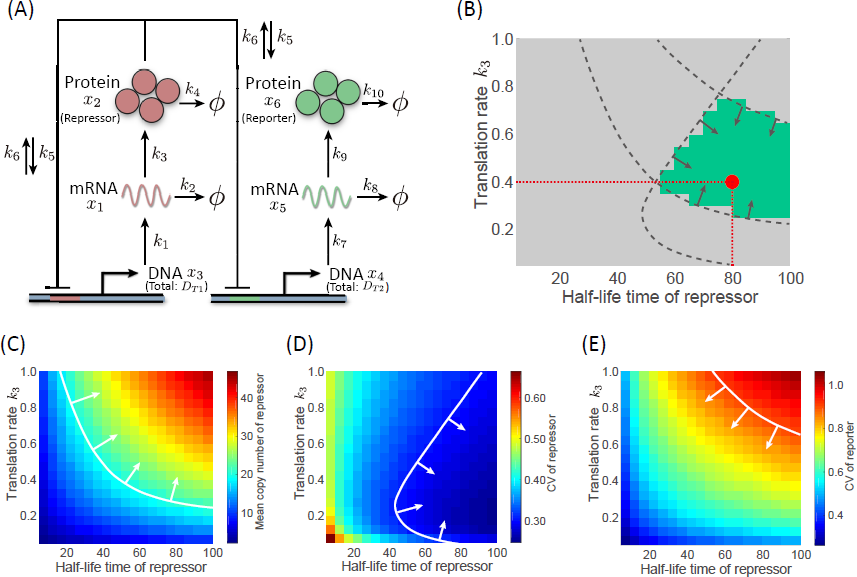
(A) Schematic diagram of a negative feedback biocircuit with a downstream reporter protein. (B) Parameter region satisfying the three design specifications. (C) The lower bound of the mean copy number of the repressor protein. (D) The upper bound of the CV of the repressor protein. (E) The upper bound of the CV of the reporter protein.

To demonstrate the optimization based synthesis method, we consider to design a bio-circuit in Fig. 4A, where a reporter protein is added to the downstream of the negative feedback circuit in Fig. 3A. As design specifications, we require the biocircuit to satisfy the following three performance criteria at steady state: (i) the mean copy number of the repressor molecule is at least 20, (ii) the CV of the repressor protein is less than 0.30, and (iii) the CV of the reporter protein is less than 0.90. These specifications are translated as

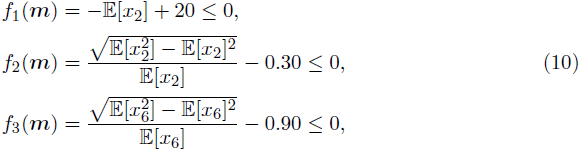

where *x*_2_ and *x*_6_ denote the copy number of the repressor and the reporter proteins, respectively.

For illustration purpose, we consider two tuning parameters, the translation rate *k*_3_ and the degradation rate *k*_4_ of the repressor protein. Note that these parameters can be tuned, for examlpe, by engineering the ribosome binding site [35] and degradation tags of the protein, respectively. Using the semidefinite programs presented in the previous section, we produced a parameter map showing the feasible design space with which the biocircuit satis es all of the three statistical design requirements in (10) (Fig. 4B). Figure 4B illustrates that the three design features specified by (10) are in a trade-off relationship in that moving a parameter to one direction satis es one constraint but violates another. Thus, we need to carefully choose parameters in the middle of the parameter space. The parameter map provides valuable information to narrow the potential combinations of genetic parts to be tested and reduces the iterations of design-build-test cycles. To verify the result of the parameter space exploration, we simulated the stochastic biomolecular reactions of the negative feedback biocircuit using the stochastic simulation algorithm [20], where the parameters were taken from the feasible design space as illustrated by the red dot in Fig. 4B. The mean copy number of the repressor protein was 26:2 copy, the CV of the repressor and the reporter proteins were 0:273 and 0:725, respectively, which all meets the design specifications.

### Noise attenuation requires balanced expression and repression

We further investigated the statistical values of the negative feedback biocircuit in detail to better understand the underlying mechanisms that limit the feasible design space in Fig. 4B and clarify design strategies (Fig. 4C–E). Figure 4C shows that increasing the translation rate of the repressor protein results in the increase of the repressor copy number at steady state despite the negative feedback. This implies that the translation of mRNA has a more in uence on the total copy number of the repressor protein than the negative feedback. Thus, a design strategy for meeting the specification *f*_1_(***m***) ≤ 0, which is to maintain the copy number of the repressor at least 20 molecules is to increase the translation rate *k*_3_. On the other hand, Fig. 4D illustrates that the CV of the repressor copy number does not decrease monotonically with *k*_3_, suggesting that the two design features *f*_1_(***m***) ≤ 0 and *f*_2_(***m***) ≤ 0 are in a trade-off relationship.

**Figure 5:**
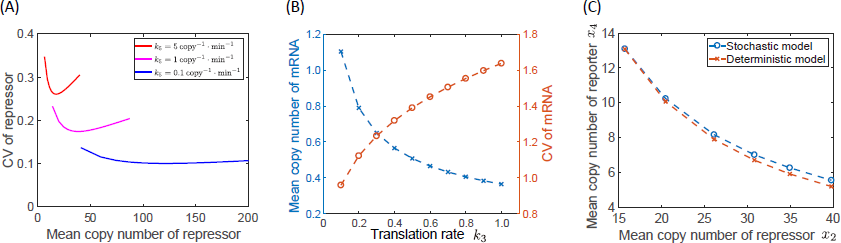
(A) Non-monotonic relation between the CV and the copy number of the repressor protein. (B) The mean and the CV of the mRNA copy number. (C) The mean and the CV of the repressor protein. The mean copy number does not follow Michaelis-Menten kinetics.

It is interesting to observe that there is an optimal strength of translation *k*_3_ that mini-mizes the CV of the repressor copy number (Fig. 4D). Since the intrinsic noise of biocircuits comes from the low copy nature of molecules, it is counterintuitive that both of the mean and the CV of the repressor increase at the same time in Fig. 4C, D. We observed that this trend is generic for a wide range of the repressor-promoter binding rate *k*_5_, which directly controls the strength of the negative feedback (Fig. 5A). We suspect that this is due to a trade-off relation between the strength of repression and transcription. More specifically, increasing the translation rate results in the attenuation of noise due to the high copy numbers of the repressor protein, but at the same time, it also increases the variance of the mRNA copy number due to the strong repression as illustrated in Fig. 5B. Then, the highly stochastic mRNA transcription indirectly contributes to increasing the dispersion of the copy number of the repressor protein. Fig. 5A suggests that the former is dominant when the translation rate *k*_3_ is small, but the latter becomes dominant as the increase of *k*_3_. These analysis results suggest that balancing the repression and the expression is a key to attenuate the noise in negative feedback biocircuits.

### Expression level of reporter protein is dependent on the structure of upstream biocircuits

A typical approach to probing protein concentrations, or internal states of biocircuits, without disrupting cells is to express a fluorescent reporter protein at the downstream of a target molecule. The internal states are then indirectly quanti ed based on the uorescence measurements. Characterizing the expression levels of the reporter versus the target molecule is thus essential for rigorously quantifying the internal states of biocircuits. For the biocircuit in Fig. 4A, the mean expression level of the reporter protein is given by 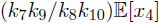 as suggested by (3), where 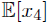 represents the mean copy number of the free DNA that is not bound by the repressor protein. To calculate 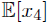, the moment equation is

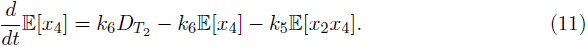

Equation (11) implies that 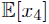 depends on the second order moment 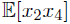, and thus, higher order moments are necessary to fully characterize 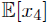. In other words, the mean copy number of the reporter 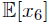 cannot be determined simply from the mean copy number of the repressor 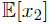 but it requires higher order statistics, which indirectly depends on moments of the upstream negative feedback pathways via moment matrices 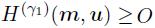 and 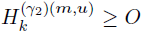 in (8). This is in contrast with the deterministic modeling, where the steady state concentration of the free DNA ***x***_4_ is expressed by the Mechaelis-Menten equation

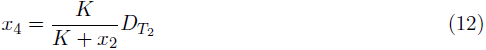

with *K* = *k*_6_/*k*_5_.

Using the convex optimization program, we characterized the reporter expression level versus the repressor copy number in Fig. 5C, where the Michaelis-Menten equation (12) is superimposed. The figure clearly illustrates that the reporter copy number deviates from the Michaelis-Menten kinetics with the maximum relative error of 6%, suggesting that the simple Michaelis-Menten kinetics is erroneous especially when the biocircuit is highly stochastic.

### Sensitivity analysis of descriptive statistics

The ability of model-based biocircuit design is currently limited by the uncertainty of parameter values in mathematical models. The source of the uncertainty partly lies in misidenti ed parameters due to insufficient and noisy measurements, but more inherently, it lies in extrinsic perturbations to host cell environments such as growth conditions. As a result, the process of biocircuit engineering often requires ad-hoc tuning of circuit parameters to deal with the deviation of circuit performance from model predictions. Sensitivity analysis allows for quantifying the impact of model uncertainties on the behavior of bio-circuits and nding sensitive design parameters that need special attention in the build process.

We developed semidefinite optimization programs to evaluate the sensitivity of the descriptive statistics in Table 1. Specifically, let 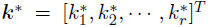 denote a vector of nominal parameters with which a biocircuit satis es performance specifications *f_i_*(***m***) ≤ 0 (*i* = 1, 2, …, *s*). We consider a parametric perturbation 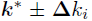, where 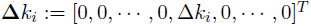 is a (small) perturbation to the nominal parameter 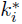. The goal of the sensitivity analysis is to find the range of the statistics under the perturbation 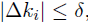, where *δ* is a given constant. This can be formulated in an optimization form as

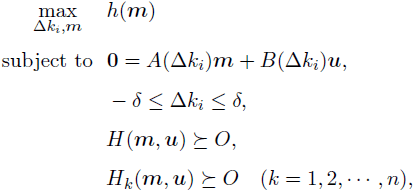

where we denote the perturbed coefficient matrices in (7) by *A*(Δ*k_i_*) and *B*(Δ*k_i_*). We convert this optimization program into a convex form to enable efficient computation of the worst-case statistics for all possible parameter combinations satisfying |Δ*k_i_*| ≤ *δ* with mathematical rigor (see Supporting Information S.6 for details). In other words, the optimization program can strictly guarantee the robustness of biocircuits for all of the parameters satisfying |Δ*k_i_*| ≤ *δ*.

Using this approach, we performed sensitivity analysis of the negative feedback biocircuit in Fig. 4A around a nominal parameter value shown as the red dot in Fig. 4B. Initially, we computed the worst-case mean and the CV of the repressor protein ***x*** when *k*_1_, *k*_2_, *k*_3_ and *k*_4_ are perturbed within 5% of the nominal value (Fig. 6). The result shows that the deviation of the statistics is almost equal between the perturbations, implying that there is no highly sensitive parameters that signi cantly affect these statistics. Figure 6 also illustrates that the mean and the CV do not violate the design constraints 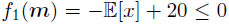 and 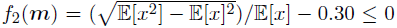, which guarantees that the negative feedback biocircuit designed in Fig. 4 is robust against these parameter perturbations.

**Figure 6:**
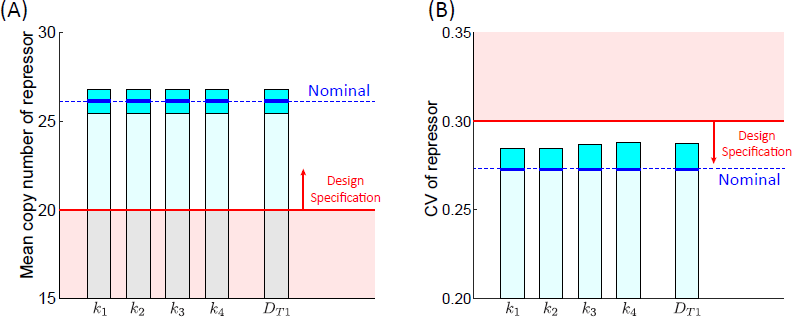
Sensitivity analysis for parameter perturbation. The red region shows the design specifications (10). (A) Sensitivity analysis of the mean copy number of the repressor protein. The blue region shows the worst-case upper and lower bounds for parameter perturbation. (B) Sensitivity analysis of the CV of the repressor protein. The blue region shows the worst-case upper bound for parameter perturbation.

Another important but often overlooked design parameter of biocircuits is the plasmid copy number, which is controlled by the replication origin of a circuit plasmid. Although the plasmid copy number is assumed constant in Fig. 4B, variance of the plasmid copy number in real biological cells affects the behavior of biocircuits. To analyze the effect of plasmid copy numbers, we applied the same approach to computing the worst-case performance of the negative feedback biocircuit against 5% deviation of the copy number of the repressor plasmid from the nominal value *D*_*T*1_. From Fig. 6, we can guarantee that the designed negative feedback biocircuit can operate within the pre-specified range of performance norms even under the extrinsic perturbation to the plasmid copy number.

## Discussion

A promising approach toward robust engineering of complex biocircuits is to guarantee the performance of individual circuit components at high precision. The highly stochastic nature of biomolecular reactions, however, hinders reliable assessment of biocircuit behaviors in current synthetic biology. To advance a model-guided design approach, it is critical to develop design-oriented theoretical tools that can rigorously certify robustness of stochastic chemical reactions.

In this paper, we have presented an optimization based approach to designing stochastic biocircuits. The presented approach allows for specifying the design features of biocircuits using intuitive and well-defined metrics of descriptive statistics. The mathematical optimization algorithms enable systematic exploration of the design space to nd parameter con gurations satisfying the specifications. In contrast with approximation based approaches [25, 36], the presented method provides mathematically rigorous certi cation of circuit performance based on user-specified statistical norms. Thus, it is amenable for robust synthesis of stochastic biocircuits that require high reliability. Moreover, the convex nature of the optimization programs allows for efficient search of the optimal solutions by bene tting from existing algorithms of mathematical optimization.

To demonstrate these features, we have explored the design space of the negative feedback biocircuit in Fig. 4A and obtained the parameter map of feasible design space with which the biocircuit satis es design requirements. In particular, the optimization based analysis elucidated that there is an optimal translation rate of the repressor protein that best attenuates intrinsic noise and that it is caused by the tradeoff relation between repression and expression. It is worth noting that a similar tradeoff relation was previously predicted for a metabolic pathway [37] and the repressilator [38] based on approximated model-based analyses. A similar U-shaped trend to Fig. 5A was also observed by experiments [39]. These examples suggest that even simple biocircuits can exhibit complex noise characteristics, which emphasizes the importance of advanced mathematical and computational frameworks for analyzing stochastic biocircuits.

Although not discussed, multi-modality of population distributions is one of the important design features that would likely to be included in the specifications of stochastic biocircuits. In synthetic biology, multimodal population distribution is often associated with multi-stability of the governing dynamics of biocircuits and is used to build switch-like systems as represented by the celebrated genetic toggle switch [15]. An optimization based approach was recently developed to design multimodal biocircuits by directly minimizing the deviation of the distribution from a desired shape [40]. As of yet, however, there has not been a well-defined statistical metric that can quantitatively certify the existence of multimodal distributions, though a recent study indicated that most information of bimodal distributions is encoded in a small number of low order moments [41]. Future work will aim to establish statistical criteria for more advanced design features to enhance rational engineering process of complex stochastic biocircuits.

## Method

### Stochastic simulations

The stochastic simulation algorithm [20] was used to simulate time trajectories of molecular copy counts. 10, 000 cells were simulated to draw a snapshot of the population distribution at 1440 minute in Fig. 2B, Fig. 3C. The following parameter values were used for the simulations. *k*_1_ = **0**.2 min^−1^, *k*_2_ = ln(2)/5 min^−1^, *k*_3_ = **0**.5 min^−1^, *k*_4_ = ln(2)/20 min^−1^ *k*_5_ = 5 copy^−1^ .min^−1^, *k*_6_ = 1 min^−1^, *k*_7_ = **0**.2 min^−1^, *k*_8_ = ln(2)/5 min^−1^, *k*_9_ = **0**.5 min^−1^, *k*_10_ = ln(2)/20 min^−1^. 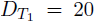 copy was used for the simulation in Fig. 2B, and 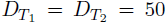 copy was used for Fig. 3C. The red dot in Fig. 4B corresponds to *k*_3_ = **0**.4 and *k*_4_ = ln(2)/80. The initial copy numbers were assumed ***x***_1_ = **0** and ***x***_2_ = **0** for Fig. 2B, and ***x***_1_ = **0** copy, ***x***_2_ = **0** copy and ***x***_3_ = **0** copy for Fig. 3B. All simulations were run by MATLAB 2016b.

### Optimization based computation of statistics

The semidefinite programs were solved with MATLAB 2016b and Sedumi 1.32 solver [42], where the following options of the solver were used. pars.eps = **0**, pars.alg = 2, pars.theta = **0**.01, pars.beta = **0**.9, pars.stepdif = 1, pars.free = 1, pars.cg.maxiter = 500, pars.cg.re ne = 10, pars.cg.stagtol = 5 10^20^, pars.cg.restol = 5 10^10^, pars.chol.canceltol = 10^20^, pars.chol.maxuden = 4000. The variables ***m*** and ***u*** were normalized as appropriate by constants to avoid numerical instability.

The truncation order *μ* was set *μ* = 8 to compute the mean and the CV of the repressor in Fig. 4C, D and, and = 6 to compute the CV of the reporter protein in Fig. 4E.

For the analysis of negative feedback biocircuits in Fig. 5, the optimization problems were solved for different values of *k*_3_. The other parameter values were set equal to those shown in the stochastic simulations section, and the truncation order was set *μ* = 8. To vary the mean copy number of the repressor at steady state in Fig. 5A, the translation rate *k*_3_ was scanned between 0.025 and 0.9. For Fig. 5C, *k*_3_ = **0**.15, 0.25, 0.40, 0.55, 0.70, 0.90 were used. The truncation order = 6 was used for the sensitivity analysis in Fig. 6.

## Acknowledgments

This work was supported in part by JSPS KAKENHI Grant Number JP16H07175 and Keio Gijuku Academic Development Funds.

